# Global decoupling of cell differentiation from cell division in early embryo development

**DOI:** 10.1101/2023.07.29.551123

**Authors:** Kalki Kukreja, Nikit Patel, Sean G Megason, Allon M Klein

## Abstract

As tissues develop, cells divide and differentiate concurrently. Conflicting evidence shows that cell division is either dispensable or required for formation of cell types. To determine the role of cell division in differentiation, we arrested the cell cycle in zebrafish embryos using two independent approaches and profiled them at single-cell resolution. We show that cell division is dispensable for differentiation of all embryonic tissues during initial cell type differentiation from early gastrulation to the end of segmentation. In the absence of cell division, differentiation slows down in some cell types, and cells exhibit global stress responses. While differentiation is robust to blocking cell division, the proportions of cells across cell states are not. This work simplifies our understanding of the role of cell division in development and showcases the utility of combining embryo-wide perturbations with single-cell RNA sequencing to uncover the role of common biological processes across multiple tissues.

## Main text

During organismal development, cell-division rates vary continuously within and between tissues to generate the correct abundance and organization of cell types (*1–4*). Cell differentiation results in changes in expression and activity of cell cycle machinery and division rate, but whether cell division reciprocally regulates cell differentiation is less clear. Studies focusing on a few specific cell state transitions have identified molecular mechanisms coupling cell cycle to signaling and transcription activity which in turn regulates cellular differentiation. These span examples where differentiation is dependent on: (1) binding of a transcription factor (TF) to its target locus during S phase, such as GATA1 to globin in erythropoiesis (*5*); (2) cell-cycle lengthening to enable accumulation of lineage-specific TFs, such as PU.1 in myeloid cells (*6*, *7*); and (3) the activity of cyclin dependent kinases that phosphorylate signaling molecules and TFs to control differentiation, such as in ES cells and neurons (*8–12*).

By contrast, prior studies in whole embryos have shown that embryos under conditions of cell cycle arrest undergo morphogenesis and give rise to cells expressing markers of some differentiated cell types including neurons and muscle (*13–17*). We do not however know whether some – or even all – cell types require cell division to correctly differentiate, as these studies necessarily focused on differentiation of a few cell types, and they only examined a few markers to evaluate differentiation for the cell types examined.

We investigated the requirement of cell division in cellular differentiation of embryos using single cell transcriptome-wide measurements as they allow unbiased and quantitative evaluation of perturbation responses in all cell types in a developing organism (*18–20*). We used zebrafish as a model of vertebrate development, focusing on a critical period between early gastrulation and late somitogenesis (6 to 24 hours post-fertilization) during which embryonic cells increase about 5-fold in number and simultaneously undergo extensive differentiation from germ layers to the formation of tens of tissues including the developing brain and spinal cord, a beating heart, skin, endothelium, and circulating blood (*18*, *20–22*) (**Fig. 1A**). This time period therefore allows critical evaluation of the requirement for cell cycle progression in the differentiation of multiple tissues from all germ layers and across several cell divisions.

**Fig. 1.**
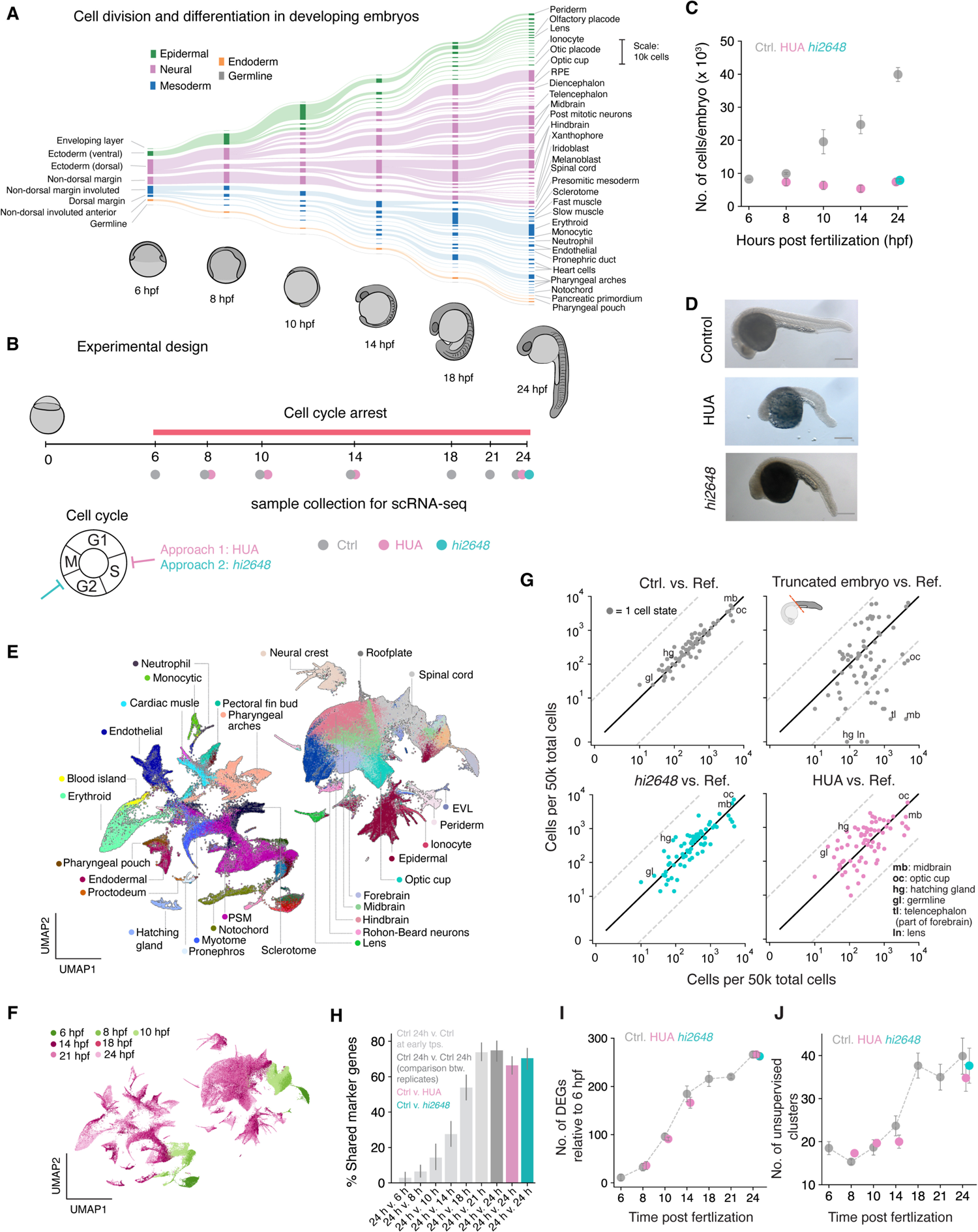
All major cell types form after cell cycle arrest. (**A**) Developmental hierarchy from 6 to 24 hours post fertilization (hpf) adapted from (*20*); flowline thickness indicates changes in cell numbers as measured by scRNA-seq. (**B**) Experimental design for cell cycle arrest: embryos developed until 6 hpf and were then arrested in cell cycle. Data was collected at specified time points between 6 and 24 hpf for control and HUA arrest and at 24 hpf for *emi1*mutant fish (*hi6248*). (**C**) Number of dissociated cells per embryo for different time points for control and arrested embryos. (**D**) Whole embryo images of control (top), HUA (middle) and *emi1* mutant (bottom) embryos at 24 hpf. Scale bar: 200 µm (**E**, **F**) UMAP representation of control data shown in (B) colored by (**E**) cell states and by (**F**) time points. (**G**) From left to right: proportion of cells of each cell type compared to reference (Wagner *et. al.* 2018) for control, truncated, *emi1* mutant, and HUA treated embryos. Solid black line represents unchanged proportions (slope=1), dashed grey lines represent 20-fold depletion/enrichment. Selected cell states are annotated; for full data see Table S2. (**H**) Mean percentage shared “marker” genes across each annotated cell state between different specified conditions (**I**) Number of differentially expressed genes between 6 hpf and further times: 8, 10, 14 and 24 hpf. (**J**) Number of unsupervised Leiden clusters at different time points. For all figures: Grey: unperturbed control embryos. Teal: *emi1* mutant. Purple: HUA treatment.

### Differentiation of all early embryonic tissues occurs in the absence of cell division

We arrested cell cycle at early gastrulation (6 hpf) using two independent approaches: a cocktail of DNA replication inhibitors HUA (hydroxyurea and aphidicolin) and a loss-of-function mutation in *emi1* (allele *hi2648*) which has been reported to arrest cell cycle at 6 hpf at the G2/M transition (*23*, *24*) (**Fig. 1B**). For HUA-treated and control embryos, we closely tracked differentiation dynamics by single-cell RNA-sequencing (scRNA-seq) between 6 and 24 hpf (**Fig. 1B**) and studied the *emi1* mutants at 24 hpf (**Fig. 1B**), for a total of 248,998 cell transcriptomes across all experiments (**Fig. S1A**). We confirmed that arrested embryos had constant cell numbers (**Fig 1C**), an increased cell size (**Fig. S1B,C**), and no DNA replication as measured by EdU incorporation (**Fig. S1D,E**). The arrested embryos continued to undergo major morphological changes as previously reported (*13–17*), including completion of epiboly, formation of anterior-posterior and dorsal-ventral axes, somitogenesis and neural tube formation. However tail elongation was severely affected (**Fig. 1D, S2**), as were cellular packing and orientation in the anterior body and head (**Fig. S1F**).

We used the time-resolved scRNA-seq data from unperturbed, HUA-treated, and *emi1* embryos to determine which cell types required cell division for their formation. To do this, we assigned all single-cell transcriptomes to annotated cell states using a classifier trained on our previously published scRNA-seq atlas of zebrafish development for all time points (*20*) (**Fig. 1E, F and Figs. S3A-F**). In control embryos, all annotated cell types were represented, and their abundances agreed quantitatively with those of the reference (**Fig. 1G**). Strikingly, we found that every one of the annotated cell states at 24 hpf were also found to be present after cell cycle arrest. This result indicates that cell cycle progression is not required for the formation of any of the cell types during the embryonic period examined.

To build confidence in this result, we carried out several controls. First, we repeated the analyses on data collected from truncated embryos whose head was removed to ensure complete loss of some cell states. We observed depletion of anterior cell states in truncated embryos as expected, establishing that we can dependably detect the presence or absence of specific cell states (**Fig. 1G**). Second, we confirmed that cells were correctly classified by examining the expression of genes specifically enriched in each cell state (defined as marker genes) in control and arrested embryos. We observed a large overlap of marker genes between cell states in control and arrested embryos at 24 hpf but not when comparing 24 hpf control cell states with earlier developmental stages (**Fig. 1H**). Third, we independently evaluated the increasing complexity of these embryos over time using both the number of genes increasing or decreasing in expression compared to the mid-gastrula (6 hpf) embryo (**Fig. 1I**) and the number of unsupervised clusters in scRNA-seq data (**Fig, 1J**). Each of these two additional metrics showed similar complexity of the wildtype and arrested embryos across the differentiation time course, without need for assignment of cell types or comparison to the reference data. These multiple metrics indicate that progression through the cell cycle is not required for differentiation during early embryogenesis.

These observations suggested that arrested embryos, which exhibit macroscopic aberrations in their morphology, are nevertheless composed of the same repertoire of cell types and states as that of normal embryos. These results were further supported by staining for specific cell types at 24 hpf using RNA fluorescent *in situ* hybridization (FISH) against cell type specific genes revealed in the data (**Fig. 2A**). The perturbed embryos contained rare cell types such as the neural tube floorplate and the pancreatic primordium, which first form during the period of cell cycle arrest and represent only 0.28% and 0.15% of cells in the 24 hpf embryo respectively as measured by scRNA-seq (*20*) (**Fig. 2B,C**). They also contained cells corresponding to progressive stages of differentiation including three blood cell types (erythroid, neutrophil precursors and monocytic cells), and two muscle types (slow and fast muscle) (**Fig. 2D,E**).

**Fig. 2.**
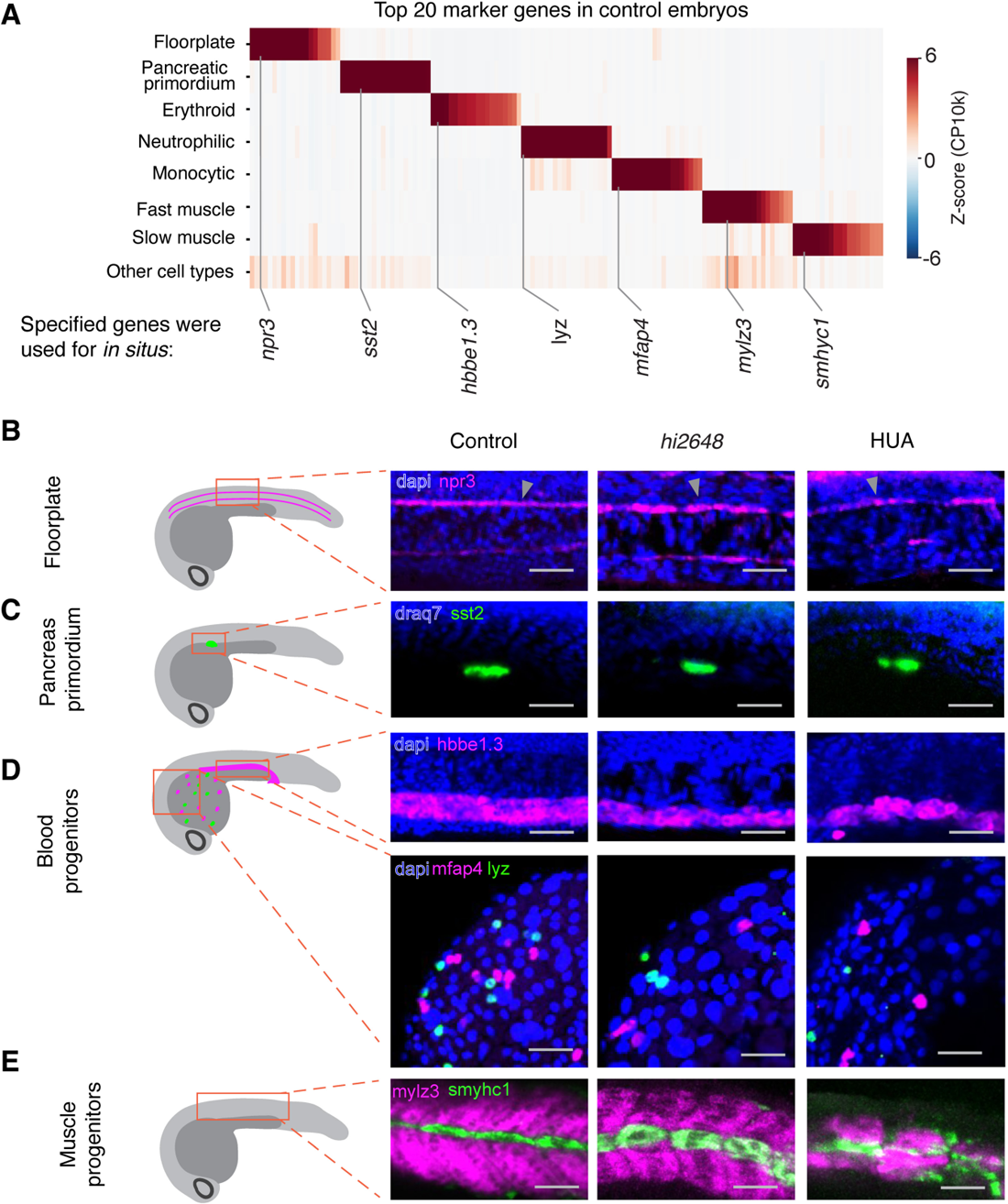
Validation of formation of cell types *in situ*. (**A**) Top 20 marker genes for select cell types (**B-E**) *In situ* hybridization of cell type specific marker genes: (**B**) *npr3* for floorplate (magenta), (**C**) *sst2* for pancreatic primordium (green), (**D**) top: *hbbe1.3* (magenta) for erythroid progenitors, bottom: *mfap4* for neutrophilic (magenta) and *lyz* for monocytic cells (green), (**E**) *mylz3* for fast muscle (magenta) and *smylc1* for slow muscle fibers (green). Scale bar: 50 µm

Together, these data independently corroborate the presence of rare and specific cell types within the perturbed embryo, and further support that all annotated cell states identified in control embryos form in the absence of cell division.

### Differentiation rates variably depend on cell division

Beyond formation of all major cell types, normal development establishes these cells in the right proportions and at the right time. Having found that all cell states can form without cell division, we next examined how differentiation timing of these cell states depend on cell division. We tested for changes in timing of differentiation by comparing single-cell transcriptomes from arrested embryos to unperturbed controls (**Fig. 3A,B**). Upon mapping each cell from 24 hpf perturbed embryos to its most similar wildtype reference (**Fig. S1A**) at any time point, we observed a delay in differentiation for both cell-cycle perturbations, with mean lags of 3.16 ± 0.02 hours in HUA-treated and 0.71 ± 0.02 hours in *emi1* mutants at 24 hpf (**Fig. 3C**), which was reproducible in all replicate experiments. Of note, this delay varies for different cell states. Erythroid cells appeared the most retarded (**Fig. 3D**,**E** 5.18 hours), recapitulating the influence of cell cycle in mammalian erythropoiesis (*5*, *25*). The optic cup, monocytic cells, presomitic mesoderm, hatching gland and other tissues showed comparable delay and the degree of lag was broadly distributed across cell types (**Fig. 3D**). We further confirmed the delay in differentiation of blood cells by quantitative PCR, and by RNA FISH: as predicted from the scRNA-seq, the erythroid globin genes *hbae3* and *hbbe1.3* accumulated more slowly upon cell-cycle arrest (**Fig. 3F,G, S4**), as did the myeloid cell markers *lyz* and *srgn* (**Fig. S4**). Therefore, although progression through the cell cycle was not required for differentiation of any cell types, its absence slowed down cell differentiation, in a tissue-specific manner.

**Fig. 3.**
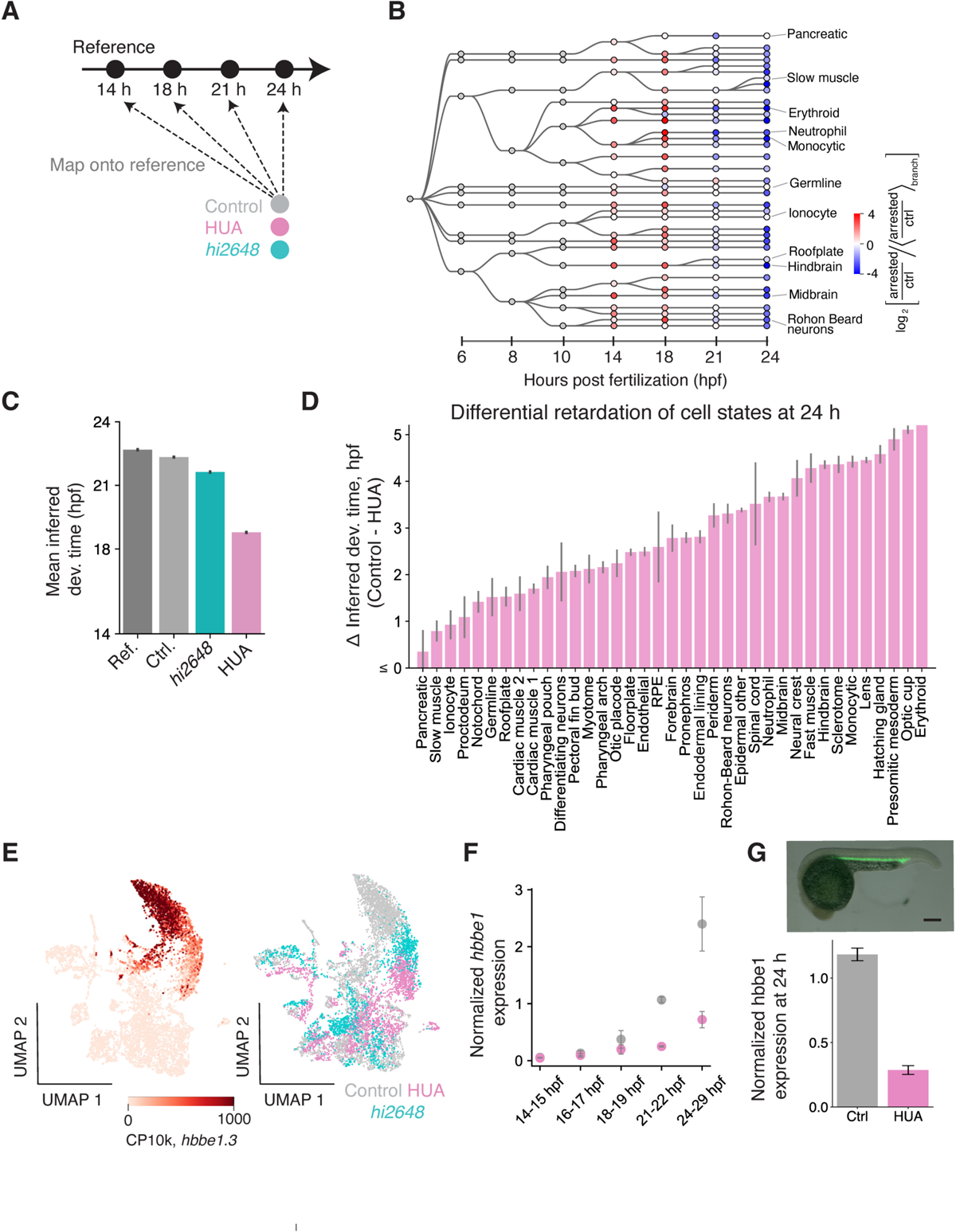
Embryonic cell types are non-uniformly retarded in differentiation upon cell cycle arrest. (**A**) Schematic for inferring developmental time of cells in the 24 hpf embryos. (**B**) Enrichment (red)/depletion (blue) of cell states across time points. It is the ratio (HUA/control) of fraction of cells in a state (shown as a circle) normalized by the geometric mean of the ratio for all cell states in the branch of the plotted developmental tree to which that state belongs. (**C**) Mean inferred developmental time across all cells for different conditions. Error bar represents SEM across cells in that condition. (**D**) Inferred delay in developmental time (Control – HUA) for all annotated cell types at 24 hpf. Error bar represents SEM across all cells in that cell type. (**E**) UMAP of all blood and endothelial cells colored by *hbbe1.3* counts per 10k total counts (CP10k) (left) and by conditions – control, HUA and *emi1* mutant cells (right) at 24 hpf. (**F**) Expression of *hbbe1.3* normalized by *gata1a* measured using whole embryo RT-qPCR at time points between 14 and 29 hpf (**G**) *in situ* hybridization of *hbbe1.3* (top) quantified in control and HUA embryos (bottom). Normalized by *gata1a* expression.

### Cell cycle arrest induces pan-embryonic transcriptional responses

Along with a delay in differentiation, we observed differences in the transcriptomes of arrested cells, which grew progressively from 14 to 24 hpf (**Fig. 4A, B, S3G**). To assess the nature of these differences, we inferred gene expression programs (GEPs) by consensus non-negative matrix factorization (cNMF)(*26*) and evaluated their contribution to total transcriptomic variation (TTV, defined in **Fig. 4C** as the magnitude of gene expression change) between control and perturbed cells in matched states (**Fig. 4C**, **Tables S3, 4**). Strikingly, just one GEP contributed to a median 33.6% of TTV across all cell types following HUA treatment (range 0.5 – 74.3% of TTV across cell types) (**Fig. 4D, left**). This program was progressively upregulated in all cell types beyond 8 hpf (**Fig 4E, F**). Another single program explained 30% of the median TTV between 24 hpf *emi1* mutant and control embryos (range 0.1 – 75.9%) (**Fig. 4D, right**). All other programs individually explained less than 2.5% TTV (**Fig. 4D**). The presence of these single global responses suggests that transcriptomic differences are driven by universal cellular responses to each form of cell cycle arrest. Many of the top genes induced by HUA were p53 target genes (*cdkn1a*, *bbc3*, *ccng1*, *sesn1*, *ddit4*, *tp53inp1*, *btg2*, *mdm2*, *plk3*, *sesn3* and *prdx1*), as expected from a response to blocking DNA replication (**Fig. 4G, H, Table S5**). Other genes enriched in this program have not been previously associated with p53 response and and could have a role in responding to replicative stress across multiple tissues (**Table S5**). The universal response to arrest in *emi1* mutants identified a distinct program enriched in DNA replication-related genes (*mcm4*, *pcna*, *rrm2*, *mcm2*, *lig1*, *top2a*, *dut*, *nap1l1*, *dnmt1* and *kpna2*) (**Fig. 4G, H, Table S5**). This program is consistent with *emi1* loss trapping cells at a G2 checkpoint (*23*, *24*). Interestingly, both global programs were deployed at low levels across cell types in wildtype embryos, and their induction upon arrest scaled with baseline expression in each cell type (**Fig. 4E**). This suggests that different cell types might be variably primed to respond to replicative stresses.

**Fig. 4.**
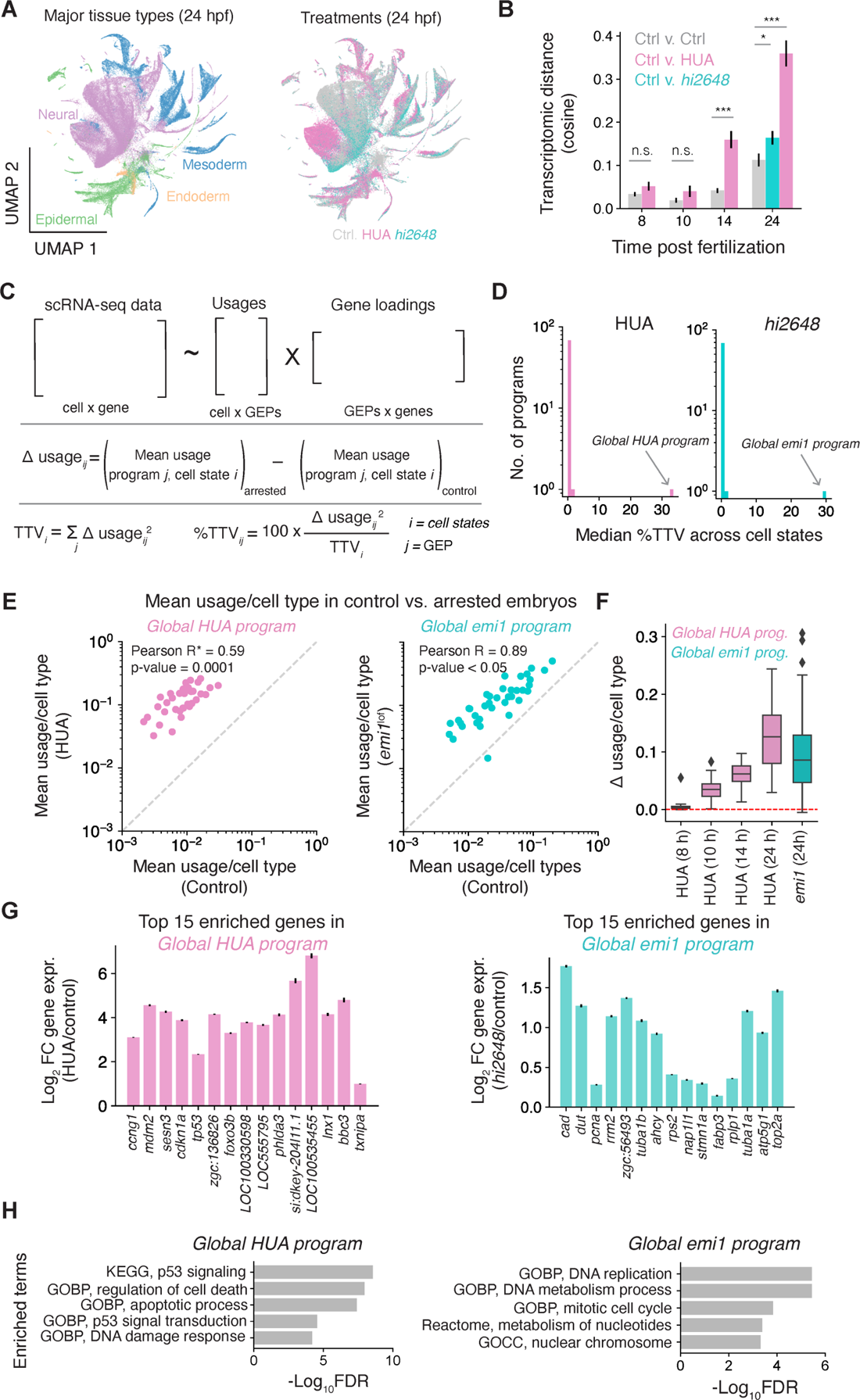
Global cell cycle arrest response across cell types. (**A**) UMAP representation of reference data colored by cell types (left) and treatment (right). (**B**) Average transcriptomic distance across cell types between different conditions and time points. Grey: control vs. control (across replicates), purple: HUA vs. control, teal: *emi1* mutant vs. control. (**C**) Schematic for finding gene expression programs (GEP) and their usages, and calculating differential (Δ) usage, total transcriptional variation (TTV) per cell type, and %TTV contributed per program (**D**) Distribution of median %TTV across cell types of all GEPs in HUA (left) and *emi1* mutant (right) embryos at 24 hpf. (**E**) Usage of HUA enriched and *emi1* enriched global programs in HUA vs. control (left) and *emi1* mutants vs. control embryos (right) respectively. (**F**) Δ usage of HUA-enriched and *emi1*-enriched program for cell states at different time points. (**G**) Fold change expression of top 15 genes of HUA enriched program (left) and *emi1* enriched program (right) between control and HUA (purple) and control and *emi1* mutants (teal) respectively. (**H**) Gene set enrichment analysis for global HUA and *emi1* mutant specific program genes. GOBP: Gene ontology biological processes; GOCC: Gene ontology cellular component.

### Cell type proportions are not robust to loss of cell division

Finally, we asked whether the proportions of different cell types in an embryo are sensitive to cell cycle arrest. Within eight hours of arrest (14 hpf), 6 out of 24 annotated cell states already showed significant changes in proportion, and by 24 hpf the number had increased to 24 out of 37 cell states (**Fig. 5A, Table S6**). These changes occurred reproducibly between the two methods of cell cycle arrest (**Fig. S5A**). These differences in proportions could be due to non-uniform proliferation rates across the embryo, because cell types emerging from highly-proliferative progenitors should be underrepresented after cell cycle arrest while those that differentiate from less-proliferative progenitors should be relatively enriched. To test this hypothesis, we estimated the expected proliferation of cells giving rise to each cell state from 6 hours to 24 hours using changes in measured cell abundances over time in a closely resolved scRNA-seq time series of untreated embryos from (*20*) (**Fig. 5B, Table S7**). The inferred rates indeed correlate with observed changes in cell-type abundance (**Fig. 5C, Fig. S5B**) (Pearson *r* = −0.47, p-value = 0.003), consistent with changes in cell type proportion resulting from loss of proliferation. However, these predicted changes over-estimate the actual changes observed by more than five-fold (**Fig. 5C**), so it is possible that some compensatory changes occur to maintain embryonic patterning upon cell cycle arrest.

**Fig. 5.**
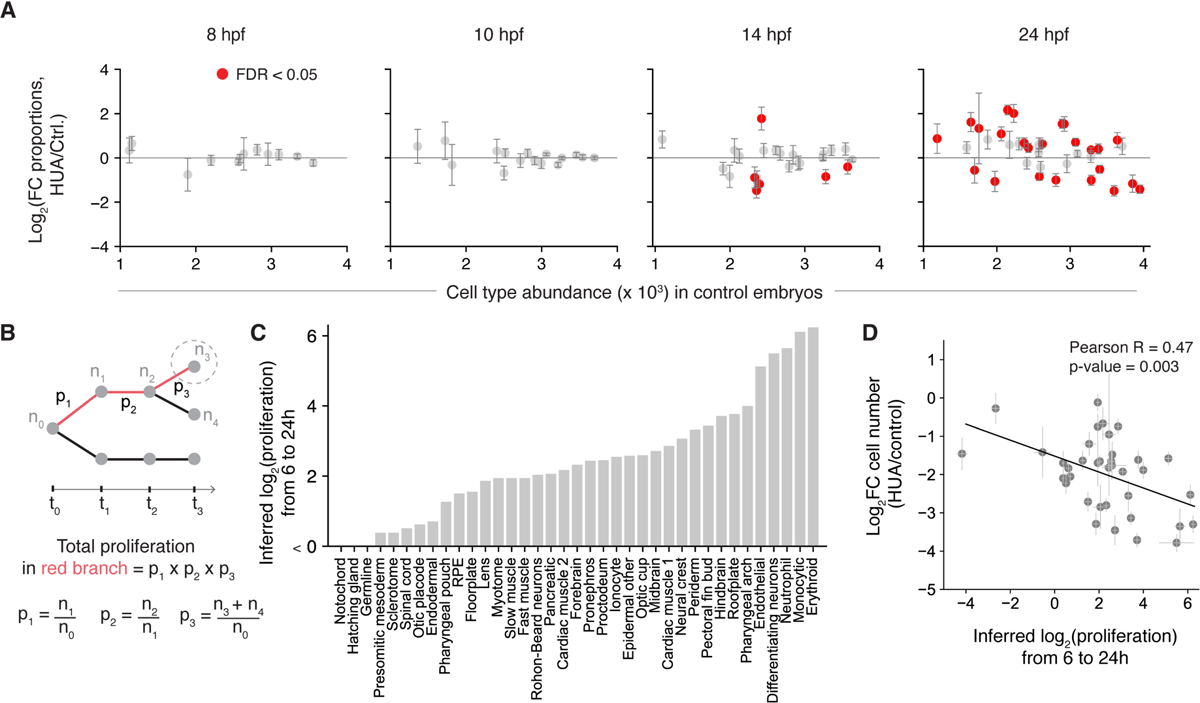
Cell type proportions are not robust to cell cycle arrest. (**A**) Fold change in proportions of cells for each annotated cell state across different time points (8, 10, 14 and 24 hpf) in HUA treated versus control samples. Significant changes are marked in red (FDR < 0.05). using beta binomial test and Benjamini Hochberg correction (**B**) Schematic for calculating the expected cumulative number of cell divisions of cells over time. p1, p2 and p3 represent proliferation between adjoining states and n0 to n4 are number of cells in labeled states calculated from the fraction of cells occupying each state as seen by scRNA-seq, and the total embryo cell counts from Ref. (*20*) (**C**) Inferred log2(proliferation) of different cell types from 6 to 24 hpf (**D**) Comparison between log2 fold change in numbers of cells in each annotated cell type versus their proliferation from 6 to 24 hpf. Black line is the trendline with slope = −0.22 Pearson correlation = −0.47.

Taken together, our results indicate that cell differentiation is not dependent on cell cycle progression, but that the proportions and total numbers of different cell types may be tuned by differential rates of cell cycle progression. We speculate that decoupling of control of cell state transitions from cell division may be a general principle that allows developmental systems to more easily evolve either changes in cell type proportion or emergence of new cell types. In light of these results, we interpret prior findings supporting the molecular control of differentiation by the cell cycle as likely reflecting specialized tissue adaptations rather than a general mechanism of development. Some of these results may reflect weak interplay between cell cycle and differentiation that quantitatively tune development processes, including affecting the rate of cell differentiation as seen here in response to cell cycle arrest. Our analytical framework makes use of an annotated, wild-type embryonic developmental atlas to quantitatively and systematically evaluate perturbations simultaneously across multiple tissues and should prove useful for studying multi-tissue responses to other perturbations.

## Data and materials availability

Raw RNA sequencing data is being made available on GEO, accession number to be provided shortly.

## Author contributions

The study was conceptualized by KK and AMK; perturbation experiments and analysis were carried out by KK; embryo truncation experiment by NP; study supervised by SGM and AMK; writing by KK, AMK.

## Acknowledgments

This work was funded by NIH grant 1R01HD096755. We thank members of the Klein lab - Sean E. McGeary, Tal Scully, James C. Taggart, Hailey M. Cambra, Laura E. Bagamery and of Megason lab - Nagarajan Nandagopal for their critical reading of the manuscript; Anna Philpott, Andrew Murray and Connie Cepko for suggestions on experimental design and analyses. We would also like to acknowledge Lexie O’Brien for helping with zebrafish maintenance and *in situ* hybridization experiments and Andrew Ross Murphy for animal care.

## Supplementary Information

### Materials and Methods

#### Zebrafish embryos and staging

AB wild-type strains were used for all HUA perturbation experiments. Zebrafish mutant line *hi2648*, a mutant for the gene *emi1* (*17*, *24*, *27*), was kindly gifted by Dr. Jennifer Rhodes. Embryos were developed within the ambient temperature of Zebrafish development. Time of fertilization at 28.5 C was used to stage each clutch and this was confirmed using morphological features of the embryos (*21*). All zebrafish were housed in a facility overseen by the Harvard Medical Area Standing Committee on Animals (our IACUC) which performs regular inspections and under which we have an approved protocol for all animal procedures.

#### Cell cycle arrest

Zebrafish embryos were arrested for cell cycle using two complementary approaches. S phase cell cycle arrest was induced using a cocktail of drugs – hydroxyurea (Sigma Aldrich H8627-5G) at 20 mM and aphidicolin (Sigma Aldrich A0781-5MG) at 150 µM concentration in 1% dimethyl sulfoxide (DMSO) (*28*) in egg water. G2/M phase arrest was induced by crossing heterozygous *emi1*^wt/mut^ zebrafish to generate homozygous *emi1*^mut/mut^ embryos, referred as *hi2648* in the manuscript. Homozygous mutants with cell cycle arrest were screened at 24 hpf as they have very clear phenotype by this time – these embryos are smaller and have bent tails and the heterozygous mutants and wildtype are indistinguishable.

#### EdU pulse labeling

Embryos were incubated in 1mL EdU (5-Ethynyl-2’-deoxyuridine) (Cayman Chemical CAS number 61135-33-9) solution for 30 minutes on a shaker at 4°C. The EdU solution contained 1X E3 media, 15% v/v DMSO, and 500 µM EdU for control and emi1 mutant embryos, and additionally 20mM hydroxyurea and 150 µM aphidicolin for S-phase blocked embryos. Embryos were then recovered at 28.5°C for 10 minutes in control media (1% DMSO in egg water) for unperturbed control embryos and emi1 mutants, and HUA solution (20mM hydroxyurea, 150 µM aphidicolin, 1% DMSO in egg water) for S-phase arrest samples. Samples were then fixed in 4% w/v PFA either for 2 hours at room temperature or overnight at 4°C. Embryos were washed twice with 1X PBS solution. They were then incubated in 1% v/v triton/PBS solution for 1 hour at room temperature on a shaker followed by a 5-minute rinse in 2% w/v BSA/PBS solution.

Embryos were then stained using Click-iT EdU cell proliferation Alexa 488 kit (Thermofisher Scientific C10425). They were then incubated in DAPI (1:10,000) or HOECHST (1:1000) solution for half hour for nuclear staining. They were washed twice with 1XPBS and either imaged right away or stored at 4°C for up to a week before imaging. Imaging was done using LSM710 upright confocal microscope using 20X water objective. A z-stack of the embryo trunk was captured.

#### EdU quantification

Three z-slices spanning across 3 different depths of the embryo trunk were used for EdU analysis. Image J’s smoothing filter (https://imagejdocu.tudor.lu/gui/process/smooth) and background subtraction (rolling ball radius of 50 pixels) was applied to every slice. DAPI channel image was binarized to obtain DAPI positive pixels (threshold chosen by eyeballing). EdU intensity of DAPI positive pixels was normalized by total DAPI positive pixel count. **Fig. S1E** shows histograms of normalized EdU intensities of DAPI positive pixels plotted for images of control and HUA treated embryos. To compare between different experiments, EdU intensities were normalized by median foreground intensities.

For cell cycle arrest assessment using EdU staining of *hi2648*, n=9 (3 embryos x 3 z-slices) for the mutant and n=12 (4 embryos x 3 slices) for the control embryos used in that experiment. For EdU staining of HUA, n=6 (2 × 3 images) for the mutant and n=12 (4 × 3 images) for control associated to that experiment.

#### Cell size quantification

Embryos were stained with DAPI (1:10,000) and Alexa Fluor 568 Phalloidin (Thermo Fisher Scientific A12380 to mark the cell membrane. Images were collected at 20X magnification on confocal microscope LSM710. For cell arrest at 6 hpf, area of periderm cells on the yolk of 24 hpf embryo was measured. Cells in the peridermal layer were identified by being the most superficial cells in the image (close to the objective) and based on their squamous morphology. Using polygon selection tool in ImageJ, cells were manually traced, and their areas recorded.

#### Embryonic dissociation for inDrops

Zebrafish embryos were dechorionated using 1mg/mL Pronase (Sigma P5147-1G) for 5-7 minutes followed by washing in egg water. Embryos were dissociated as previously described in Wagner *et. al.* 2018 (*20*). Briefly, embryos were dissociated in 0.5mL LoBind microcentrifuge tubes that had been precoated with 10% w/v BSA for about 15 minutes at room temperature.

They were homogenized in 1XDPBS/1% w/v BSA for 6 to 10 hpf embryos (early time points) and in 500 µL FACSmax cell dissociation solution (Genlantis T200100) for 14 to 24 hpf embryos (late time points). Cells were then filtered through a 40µm cell strainer mesh (Fisher 352340) and centrifuged in a swinging bucket rotor at 200g for 1 minute (early time points) or 300g for 5 minutes (late time points). Cells were then washed in 1XDPBS/1%BSA using the same centrifuge settings. They were then resuspended in 1XDPBS/0.5%BSA/18% optiprep density medium (Sigma D1556-250ML). Cell concentration was manually quantified using hemocytometer and adjusted to a final concentration of ∼100,000 cells/mL before encapsulation.

#### Single-cell Sequencing and Counts matrix

Single-cell transcriptomes were barcoded using inDrops (*29*, *30*). Standard transcriptome RNA-seq libraries were processed as reported in Zilionis *et. al.* using inDrops v3 protocol. The transcriptome libraries were sequenced on reads Illumina NextSeq 500. Libraries used standard Illumina sequencing primers and 61 cycles for Read1, 14 cycles for Read2, 8 cycles each for IndexRead1 and IndexRead2. Raw fastq files was processed using inDrops.py pipeline (github.com/indrops/indrops). Sequenced reads were mapped to a zebrafish reference transcriptome built from the zebrafish GRCz10 genome assembly (Assembly Accession: GCF_000002035.5) using bowtie version 1.1.1 (*20*).

### *In situ* hybridization using HCR

HCR in situ hybridization was performed using zebrafish-specific protocol from molecular instruments https://www.molecularinstruments.com/ based on third generation split probe approach from Choi et. al. 2018 (*31*). The reaction volumes were reduced to 150 µl instead of 500 µL mentioned and all reactions for *in situ* hybridization were conducted in 200 µL PCR tubes instead of the standard 1.5 mL microfuge tubes described in the molecular instruments protocol. The following cDNA sequences (NCBI accession number) and HCR amplifiers were used to design probes:

| S.No. | Gene | Tissue/cell type | Amplifier | Accession number |
| --- | --- | --- | --- | --- |
| 1 | *hbae3* | Erythroid | B1 | NM_183066.3 |
| 2 | *hbbe1* | Erythroid | B2 | NM_001365263.1 |
| 3 | *gata1a* | Erythroid | B3 | NM_131234.2 |
| 4 | *npr3* | Floorplate | B1 | NM_001102673.1 |
| 5 | *sst2* | Pancreatic<br>primordium | B2 | NM_131727.2 |
| 6 | *smyhc1* | Slow muscle | B1 | NM_001020507.1 |
| 7 | *mylz3* | Fast muscle | B3 | NM_131619.3 |

### Imaging and quantification of *in situ* hybridization

Images were taken at 20x magnification on confocal microscope LSM710 for Fig. 2B-D and LSM980 for Fig. 2E. For quantification of *hbbe1* at 24 hpf embryos (Fig. 3H**, Fig. S4B**), the following steps were performed using MATLAB 9.5:

1. Gaussian smoothing was performed for each of the channels – *hbbe1*, *hbae3* and *gata1a* using the function `imgaussfilt` and threshold radius =1.
2. All these genes are co-localized in the erythroid cells. Hence, we set a threshold signal from one of the genes (*hbae3*) to identify the erythroid cells in the embryo. We used Otsu method for finding a threshold using the MATLAB function `multithresh` with parameter `N = 1`.
3. We summed the intensity of signal for each of the three genes over the region of interest and divided total signal of hemoglobin genes (*hbbe1* or *hbae3*) by total signal from *gata1a* to get the values plotted in Fig. 3H and **Fig. S4B**.

### RT-qPCR experiment and analysis

Embryos were flash frozen using liquid nitrogen and stored in −80C until we collected them for all time points. RNA extraction was done using Qiagen RNAeasy mini prep kit (Catalog number: 74136). cDNA prep was done using High-Capacity cDNA Reverse Transcription Kit (Catalog number: 4368814). qPCR was performed using KAPA SYBR® FAST qPCR Master Mix (2X) (Catalog number: KK4601).

The following primers were used:

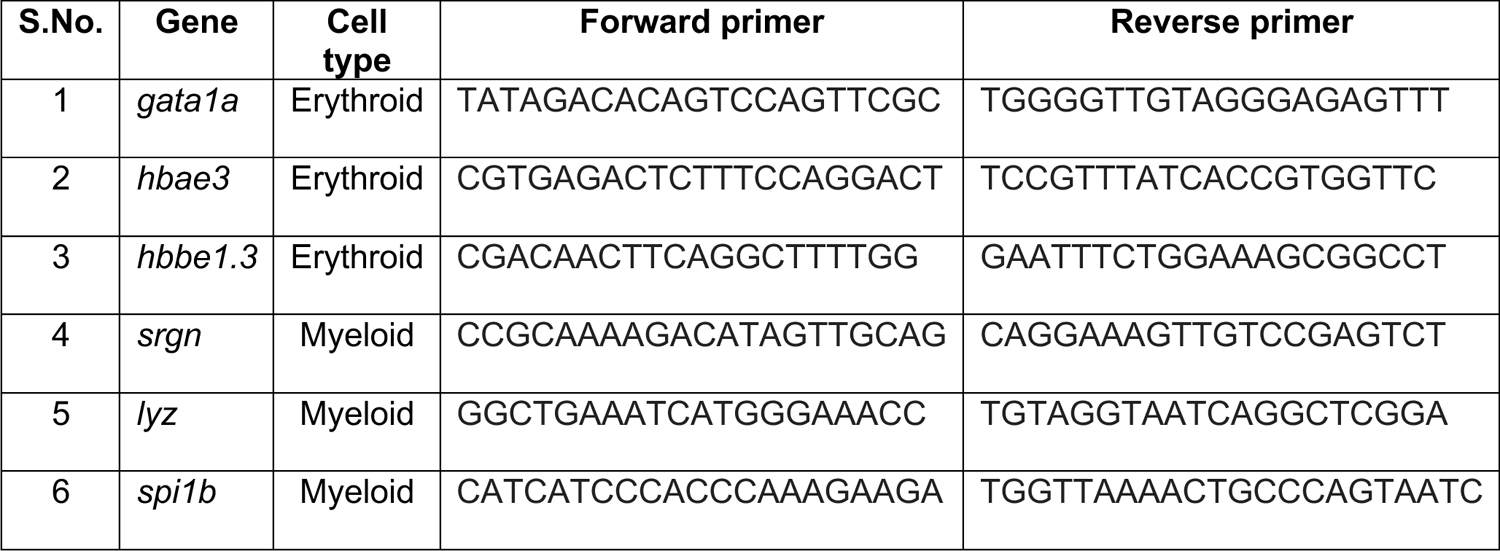

Gene expression was calculated using the standard ΔΔCt method (*32*), with normalization to 21hpf for all genes, and using *gata1a* as the reference gene for red blood cell genes *hbae3* and *hbbe1.3*, and *spi1b* as the reference gene for myeloid cell genes *lyz* and *srgn*.

### Single cell RNA-seq Data preprocessing

Scanpy (version: 1.9.1) was used for data processing steps unless otherwise stated.

1. *Cell Filtering*: inDrops data was filtered to only include UMIs from abundant cell barcodes. Transcriptomes with more than *N* total UMI counts were retained, and this threshold was chosen separately for each library after manual inspection of the UMI/cell distribution per library. The final threshold values used can be determined from the minimal total UMI counts/cell per library in metadata file `meta_data_unfiltered.tsv` on GEO.
2. *Total count normalization*: UMI counts were normalized to 10,000 total counts per cell. We used `scanpy.pp.normalize_per_cell` function.
3. *Gene filtering*: Genes expressed (count > 0) in less that 3 cells were filtered out using `scanpy.pp.filter_genes` function. We then filtered out genes with low dispersion using the scanpy’s function `scanpy.filter_gene_dispersion` with the parameters `min_mean=0.01`, `max_mean=3` and `min_disp=0.5`. A list of cell cycle and housekeeping genes were filtered out to remove confounding features for downstream analysis (*20*). A “growing” list of genes was generated by finding genes highly correlated (coefficient of correlation = 0.3) to a predefined list of cell cycle and housekeeping genes. This list was further grown by finding genes correlated to the new list generated. These genes identified by two rounds of correlation analysis were then discarded for downstream analysis.
4. *Scaling data*: Z-scoring on was performed for each gene using using `scanpy.pp.scale` function.
5. *Dimensionality reduction*: Dimensionality reduction was performed using principal component analysis using scanpy function `sc.tl.pca` with number of principal components = 50.

### Classification of cells

To classify cells, a logistic regression classifier was trained on annotations of reference data from Wagner et. al. 2018. Python’s machine learning package sklearn (version: 1.2.0) was used with function and parameters: `sklearn.linear_model.LogisticRegression(c_param=1, n_iterations = 1000)`. These states were used for Fig. 1G. For all downstream analyses such as comparing marker genes, calculating differentiation delay, and differences in cell states proportions, these cells states were renamed to more interpretable annotations and coarse grained into larger states to increase statistical power of the analyses (**Table S1**). **Figs. 1E and H, 3, 5** and **S3** contain these updated annotations. After coarse-graining, we performed knn-smoothing, that is, for every cell, we found 10 nearest neighbors in its 50-dimensional PC space and that cell was reassigned the state of maximum number of its neighbors. Python sklearn function `sklearn.neighbors` was used to find nearest neighbors of a cell. Classification was done separately for each time point and for the two experiments (**Fig. S1a**). Since Wagner et. al. data did not contain 21 hpf, we used the 24 hpf data for its classification because 21 hpf embryo looks closest to 24 hpf embryo amongst the time points in the reference data.

### Single cell RNA-Seq data visualization using UMAP

For all the UMAPs in the paper, we performed the above-mentioned scRNA-seq data processing steps, constructed a nearest neighbor graph and then plotted UMAP using this graph as input. For Fig. 1E**,F**, we constructed UMAP embeddings for control mesoendoderm and germline cells, and separately for control neuroectodermal cells. We used scanpy’s function `sc.pp.neighbors` with the parameter `n_neighbors = 100` to find nearest neighbors and the function `sc.tl.umap` with parameter `spread = 2.5` to construct the UMAP. For Fig. 3D, we constructed a UMAP embedding using blood and endothelial cells (precursors of hematopoietic cells) from 14 and 24 hpf of the perturbation experiment. We used the parameters `n_neighbors = 20` for constructing k-nearest neighbor graph and `spread = 2` for the UMAP. For **Fig. S3**, we constructed UMAP embeddings for cells from each individual time point and experimental condition. We used batch balanced k-nearest neighbors or bbknn to perform batch correction (*33*) using python function `bbknn.bbknn` version 1.51. Following parameters were used: In **Fig. S3A-C**, `neighbors_within_batch = 5` for bbknn and `spread = 2` for UMAP for all time points; in **Fig. S3D-G**, `neighbors_within_batch = 3` for bbknn and `spread = 1` for UMAP for 6, 8, 10 and 14 hpf and `neighbors_within_batch = 8` for bbknn and `min_dist = 0.4`, `spread =4` for UMAP.

### Shared marker genes analysis

Marker genes were defined as previously in Zilionis et. al. 2019 (*34*). Following this paper, we define *cell subset marker genes* as follows: Let *S* be the set of all cells considered, partitioned into *n* disjoint subsets *s*_1_*,…, s*_n_. Define *g*^-^_ij_ as the average expression (CP10k) of gene *j* in subset *i*, and *O*^-^_i_^(j)^ as the values of *g*^-^_ij_ for gene *j* sorted in descending order. Define 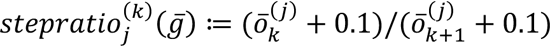 as the ratio of the k-th highest average subset expression of gene *j* to the (*k*+1)-th expression value, after adding a pseudo value of 0.1 to all average gene expression values.

A gene *j* is a marker gene for cell subset *s*_i_ if:

1. Gene *j* is detected in at 0.01 fraction of cells in *S*.
2. Gene *j* is statistically significantly higher in expression in subset *s*_i_ compared to the complement set (all cells *S* not in *s*_i_). To establish significance, we used a two-tailed Mann-Whitney U test with multiple hypothesis correction, Benjamini Hochberg FDR<5%.
3. Gene *j* has maximal average expression in subset *i*, i.e. *g*^-^_ij_ = *O*^-^_1_^(j)^.
4. Gene *j* satisfies *stepratio*_j_^(1)^(*g*^-^) > 1.2, i.e., the max-to-second-max ratio is at least 1.2x.

#### Comparing marker genes across conditions

Let the two sets of marker genes of cell subset *s*_#_ and *s*_’_ for conditions *a* and *b* be *J*_ia_ and *J*_ib_ such that *J*_ia_ = {*j* ∈ marker genes of *s*_i_ for condition *a*} and *J*_#*_ = {*j* ∈ marker genes of *s*_k_ for condition *b*}. Then, fraction shared genes between the two sets is defined as 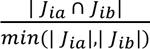

For Fig. 1H, the following comparisons have been made from left to right: Pairwise comparison between different cell states for 24 hpf control data, comparison between cell state at 24hpf and the corresponding cell state at earlier time points (6, 8, 10, 14, 18 and 21 hpf) that gave rise to the state at 24 hpf, comparison between 4 replicates of 24 hpf control data with the other 4 replicates (there are a total of 8 control replicates – 5 for HUA and 3 for *emi1* perturbation), and comparison between 24hpf control states with 24hpf HUA and 24 hpf *emi1*.

### Quantifying embryo complexity

Embryo complexity was quantified using the following metrics:

#### Number of differentially expressed genes from 6 hpf

Tests for differential gene expression were carried out between multiple sets of conditions as shown in Fig. 1I. For comparisons between 6-18hpf and 6-21hpf, we used data from reference experiment for these time points (**Fig. S1a**). All other comparisons used data from perturbation experiment (**Fig. S1a**). Genes identified as significantly different at 5% false discovery rate using a two-tailed Mann-Whitney U test with the Benjamini-Hochberg multiple hypothesis correction were considered as differentially expressed. A pseudo count of 1 CP10k was added before calculating fold-changes. Fig. 1I shows mean differentially expressed genes across comparisons. The error bars represent standard error of mean.

#### Number of Leiden clusters

For each replicate and condition, data was processed using scanpy as described in the section on “scRNA-seq data preprocessing” from steps 2 through 5. This was followed by finding 5-nearest neighbors of the data using `sc.pp.neighbors(adata, n_neighbors=5, use_rep=’X_pca’)`and then finding Leiden clusters using `sc.tl.leiden(adata, random_state = 0)`.The mean number of Leiden error across all replicates in a particular condition and time point were plotted (Fig. 1J). Error bar represents standard error of mean.

Similar to above, 18 and 21 hpf data is taken from reference experiment while other time points are from the perturbation experiment (**Fig. S1A**).

### Inferring developmental time

To infer developmental time of 24 hpf control and perturbed data (Fig. 3A**-D**), we classified cells as described in the section “Classification of cells”, though this time, we used the training data that consisted of multiple time points (14,18, 21 and 24hpf) from high time resolution wildtype reference (Reference experiment as in **Fig. S1A**). Hence, now for every condition, *r*, each cell *i*, is assigned a label *t*, *s* where *t* is the time and *s* is the cell state at that time.

Fig 3B: Let *k_t,s,p_* be the number of times a label *t,s* is assigned to a cell in the perturbed condition and *k_t,s,c_* be the number of times that label is assigned to a cell in the control condition. Let *N_p_* and *N_c_* be the total number of cells in the perturbed and control conditions respectively.

Then we calculate the ratio (perturb/control) of the votes for assigned label as 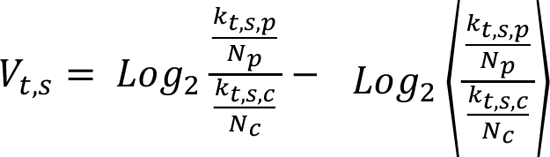, where ⟨.. ⟩ is the geometric mean across the states of the branch of the developmental tree to which that state belongs. The colors on the Fig. 3B visualizes ratios.

Fig. 3C: We calculate mean inferred time for a condition (control, HUA, *emi1*) as the average of the assigned time label *t*, 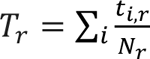 where *N*_r_ is the total number of cells in condition *r* (Fig. 3C).

Fig. 3D: To calculate delay in differentiation for each state, we group cells belonging to a particular state as previously annotated (described in the section “Classification of cells” and visualized in **Fig. S3**). Then, for every cell state in a condition, the inferred developmental time for that state is calculated as the mean of inferred time *t* of cells belonging to that state.

### Transcriptomic distance between samples

The transcriptomic distances plotted in Fig. 4B show a cosine distance between mean gene expression profiles, calculated over all highly variable genes in the transcriptome. Mean gene expression for a set of cells Ω was calculated from normalized scRNA-Seq counts matrices as 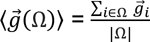. The cosine distances were calculated using `sklearn.metrics.pairwise.cosine_similarity`. Each distance *D(Ω^jk^_ab_, Ω^lm^_ab_*) is evaluated for cell sets identified at matched time points *a* and cell states *b*, for differing replicates *j* or conditions *k*. For within-condition comparisons (grey bars in the figure Fig. 4B) *k* = *m* ∀ conditions and *j* ≠ *l* ∀ replicates. Each bar is the mean of these distances across time points and cell states, and error bar represents standard error of mean. For comparisons between conditions (purple and green bars – HUA treated versus control and *emi1* mutants versus control embryos respectively), *k* = control and *m* = *emi1* mutants or HUA treated embryos and *j* = *l* are matched replicates of control and perturbed embryos from the same experimental batches.

### Gene expression programs analyses using cNMF (Fig. 4)

Gene expression programs were learnt by consensus non-negative matrix factorization (cNMF) (as described in (*26*) using the Python package for cNMF, version 1.1) with parameters: number of components k = 70, percentage of replicates used as nearest neighbors for outlier detection = 30%, and local density threshold for defining outliers = 0.2. Usage and program matrices are provided in **Tables S3 and S4**. The resulting program usages, *u*_ji_, are normalized such that ∑_j_ *u*_ji_ = 1 for programs *j* in each cell *i*. The gene weights *w*_jk_ are normalized such that ∑_k_ *w*_jk_ = 1 for program *j* over all genes *g*_k_. Two of the 70 programs (program number 38 and 8 in **Table S4**) where enriched across multiple cell types in HUA and *emi1* perturbation respectively.

Median % TTV (Fig. 4D):

For every program *j* and condition *r*, we calculated mean usage per cell state *i* as 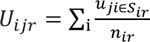 where *S*_ir_ and *n*_ir_ are respectively the set of cells from treatment *r* and cell state *i*, and *n*_ir_ = |*S*_ir_|. These values are used to plot Fig. 4E.

In Fig. 4F, we plot the difference Δ*U*_ij_= *U*_ijp_– *U*_ij0_ where *r*=0 denote the control condition. Plots show the distribution of Δ*U*_ij_ across cells states for each time point, for HUA and *emi1* conditions.

We define total transcriptional variance (TTV) per cell state, *TTV*_i_ = ∑_j_ Δ*U*_ij_^3^. We calculated the percent TTV as 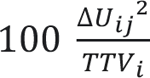, and the median %TTV usage across cell states is then 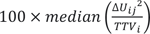. Fig. 4D plots histogram of these values for *p* = HUA and *p* = *emi1* mutant at 24 hpf.

#### Gene set enrichment analyses (Fig. 4H)

The 50 genes with highest program loadings for each global program were analyzed for gene set enrichment on the MSIGDB database and the online tool - https://www.gsea-msigdb.org/gsea/login.jsp (*37*, *38*).

### Cell type abundance quantification

Fold change in cell type abundances were assessed by calculating cell type frequencies in control and perturbed conditions. For Fig. 5A, we plot 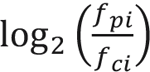 a where *f_pi_* corresponds to the frequency of cell type *i* in the perturbation condition p which could be HUA treated or *emi1* mutant embryos and *f_ci_* is the respective cell type frequency for the untreated control. Beta binomial test (*35*, *36*) and Benjamini Hochberg correction was used to evaluate statistical significance in the changes in cell type frequencies. Standard error of mean (SEM) was calculated for frequencies across replicates for different conditions. A standard error propagation formula was used to calculate SEM for fold change abundance:

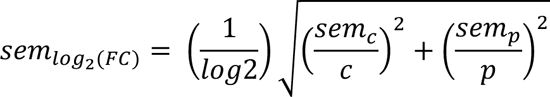

The above SEM values were used to plot error bars.

### Inference of proliferation for different cell types (Fig. 5B-D)

Inferred cell divisions from 6 hpf to 24 hpf was calculated in the following steps:

1. A development tree connecting cell states at different time points was adapted from Wagner *et. al.* 2018(*20*). The tree relationships are reported in **Table S6**.
2. We inferred total number of cells *n*_i_ in each state *s*_i_ by multiplying cell state frequencies *f*_i_ from single cell data to total number of cells *N*_1_ in the embryo at that time point. Thus *n*_i_ = *N*_1_ × *f*_i_ The total cell numbers at different time points are taken from Fig. S1C of Wagner *et. al.* 2018.
3. Proliferation between a node to an adjacent node from a time point to the next time point was calculated as follows: Let a node of a cell state *s*_i_ have a set of daughters *D*_i_ = {*s*_’_ ∈ daughters of *s*_i_}, proliferation of the cells *s*_i_ to the next time point is defined as 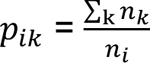
4. To find proliferation from a state *s*_j_ at time point t_?_ a state *s*_;_ at final time point t_D_: let *S*_$;_ = {*s*_#_ ∈ states connecting *s*_$_ to *s*_$_ where *l* > *j*}, *P*_jl_ = ∏^l^_j_ *p*_*ik*_ (Fig. 5B)
5. Finally, we calculated the proliferation of coarse-grained cell states by taking the mean proliferation scores of set of cell states belonging to that coarse grain cell type (**Table S1 for coarse graining of cell states**). The log_2_ of final proliferation values are plotted in Fig. 5C and compared against fold change abundances in Fig. 5D.

**Figure S1.**
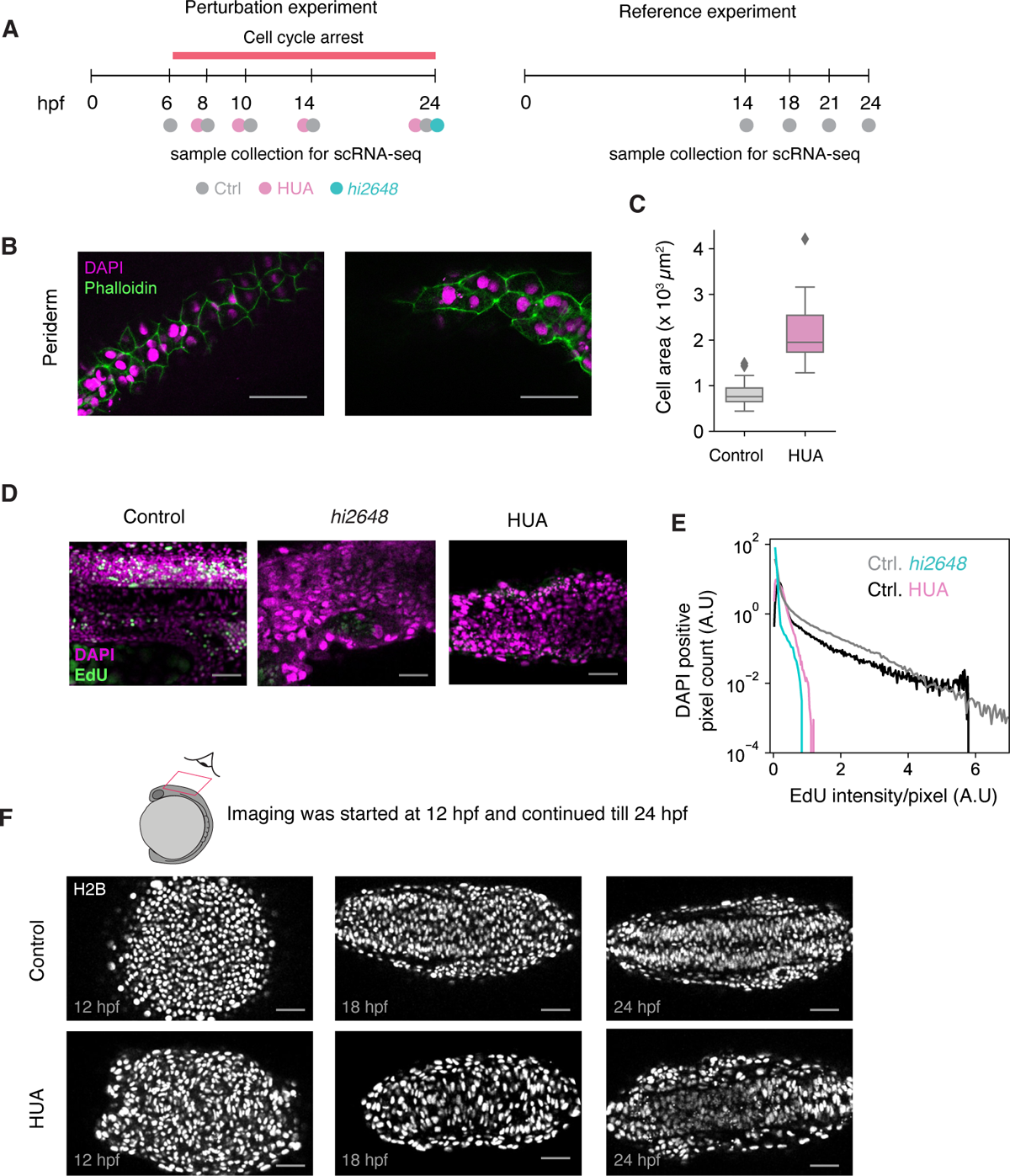
(A) Experimental design of the single cell experiments performed in the study. (B) Representative images of skin cells in control and HUA embryos. Scale bar = 50 µm. (C) Quantification of cell size in control and HUA embryos. Number of cells for area measurement: n = 52 cells (3 embryos) for control, n = 34 cells (3 embryos) for HUA embryos. (D) EdU staining in control (left), *emi1* mutant (center) and HUA treated embryos (right). Magenta: DAPI, green: EdU, Scale bar = 50 µm. (E) Quantification of EdU intensity in embryos. For EdU staining of *hi2648*, n=9 (3 embryos x 3 z-slices) for the mutant and n=12 (4 embryos x 3 slices) for the control embryos used in that experiment. For EdU staining of HUA, n=6 (2 x 3 images) for the mutant and n=12 (4 x 3 images) for control associated to that experiment (F) Representative images of dorsally mounted embryos showing neural tube folding in control and HUA treated embryos. Scale bar = 50 µm.

**Figure S2.**
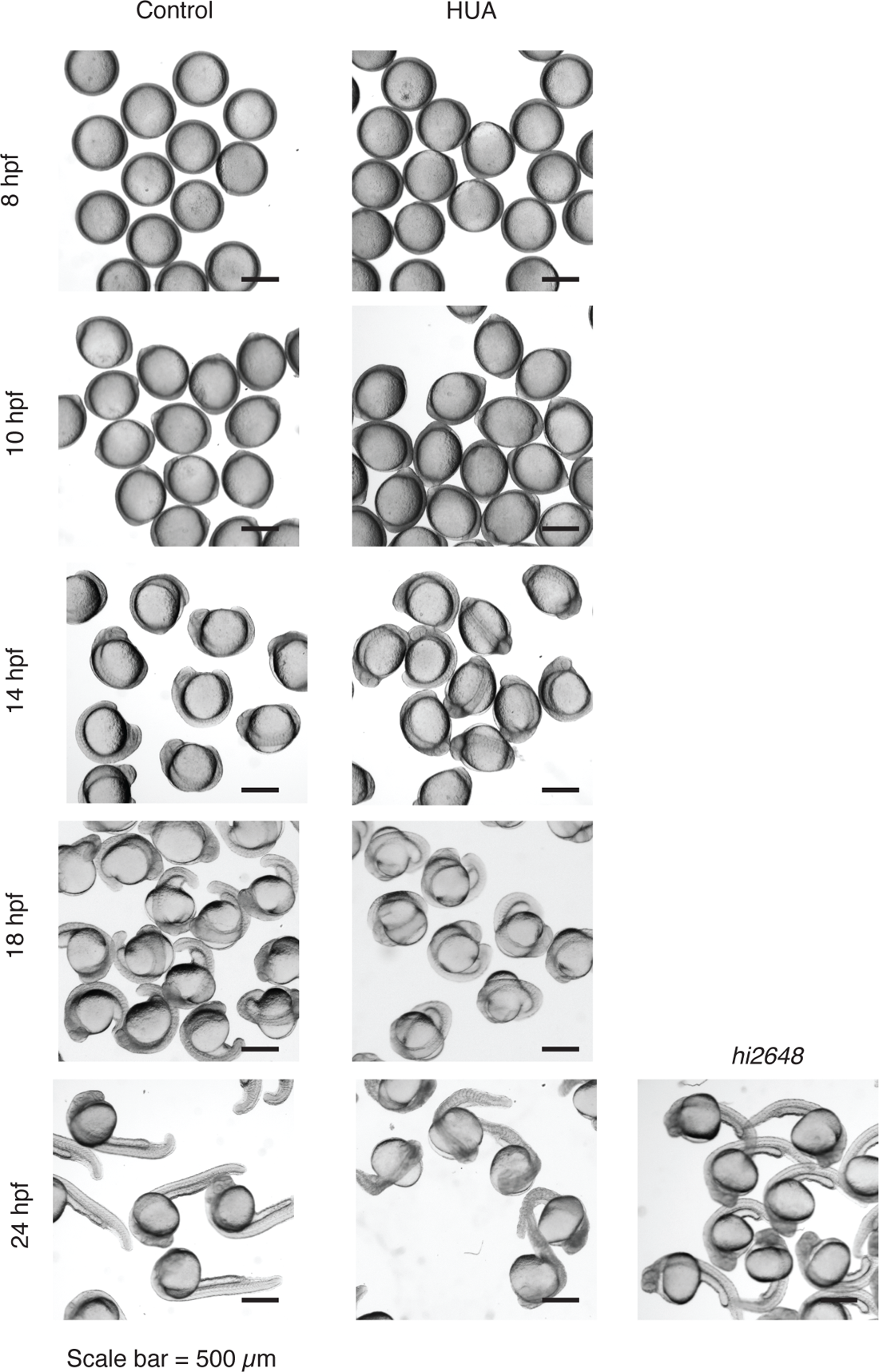
Bright-field images of control and cell cycle arrested embryos at different time points. Scale bar = 500 µm.

**Figure S3.**
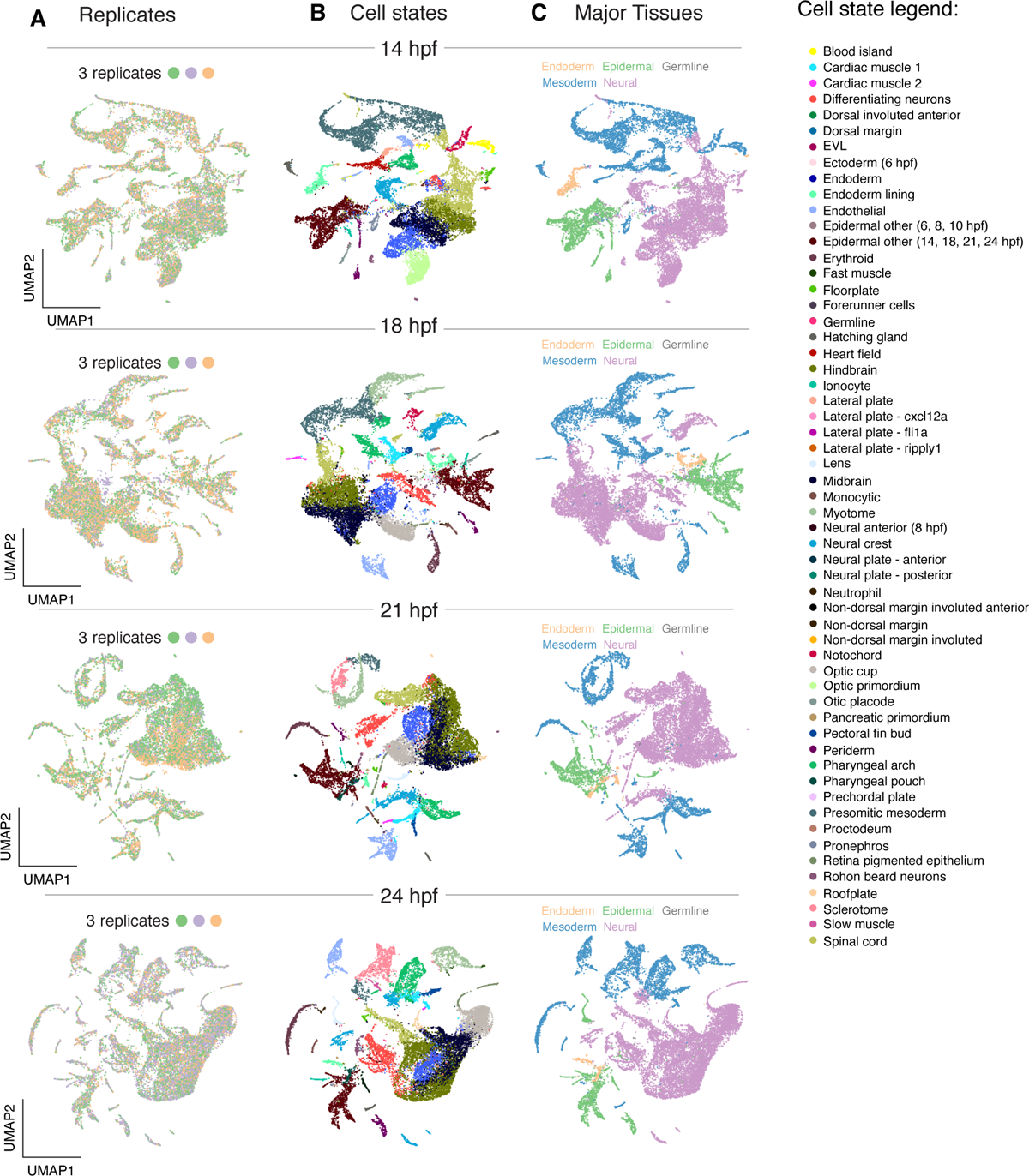
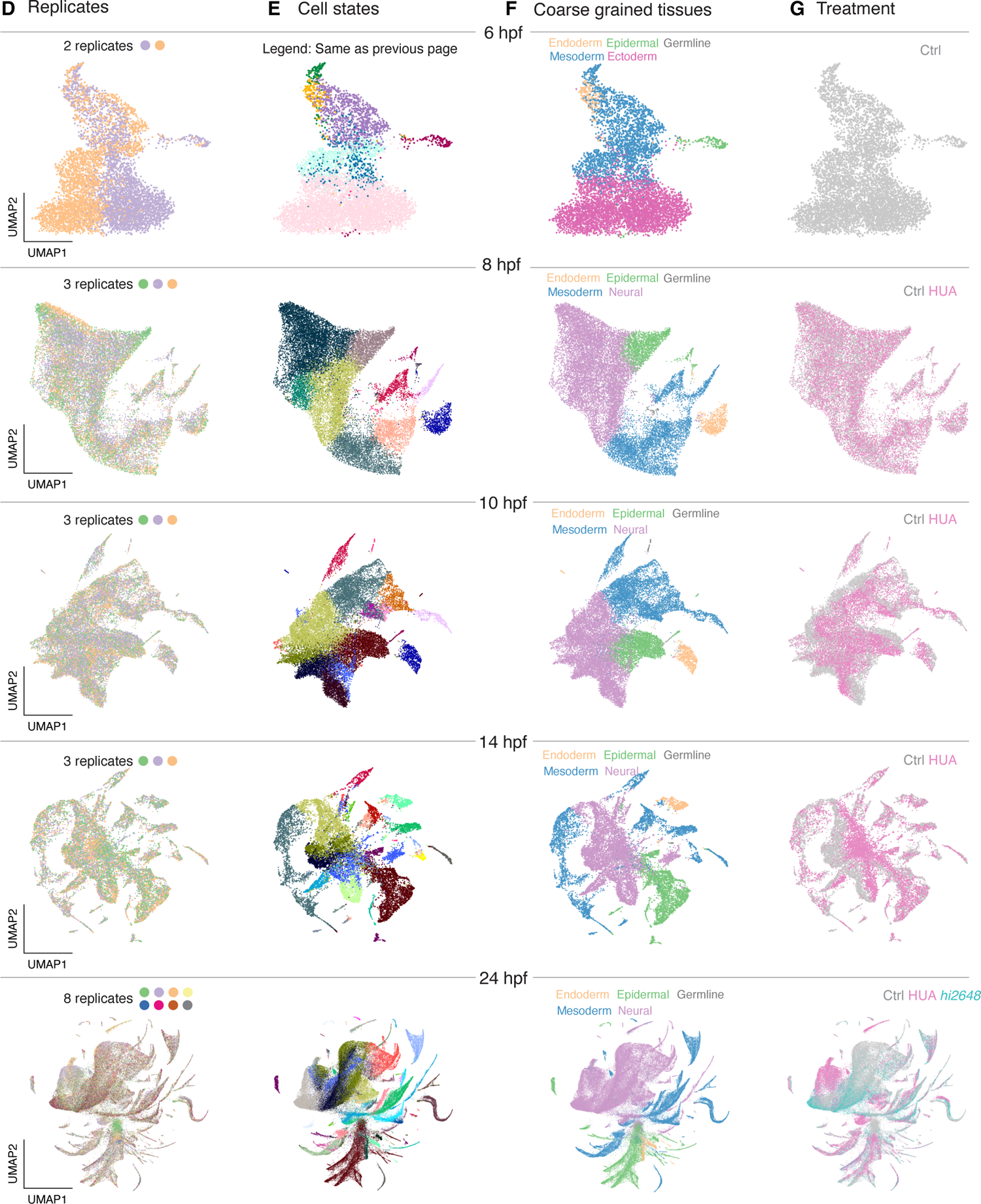
(A-C) UMAP representations of scRNA-seq data from different time points in the reference experiment and (D-G) in the perturbation experiment colored by (A,D) replicates, (B,E) cell states, (C,F) major embryonic tissues and (G) treatment.

**Figure S4.**
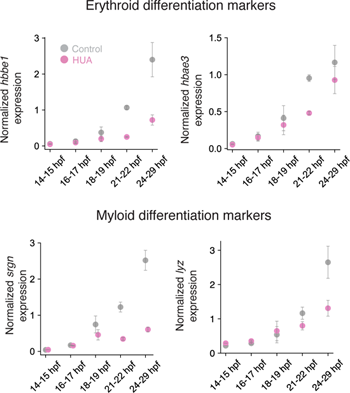
Quantification of expression of erythroid (top) and myeloid (bottom) markers of celluar differentiation across time points in development using RT-qPCR. Gene expression is normalized to cell type specific genes *gata1a* and *spi1b* respectively and to expression at 21hpf.

**Figure S5.**
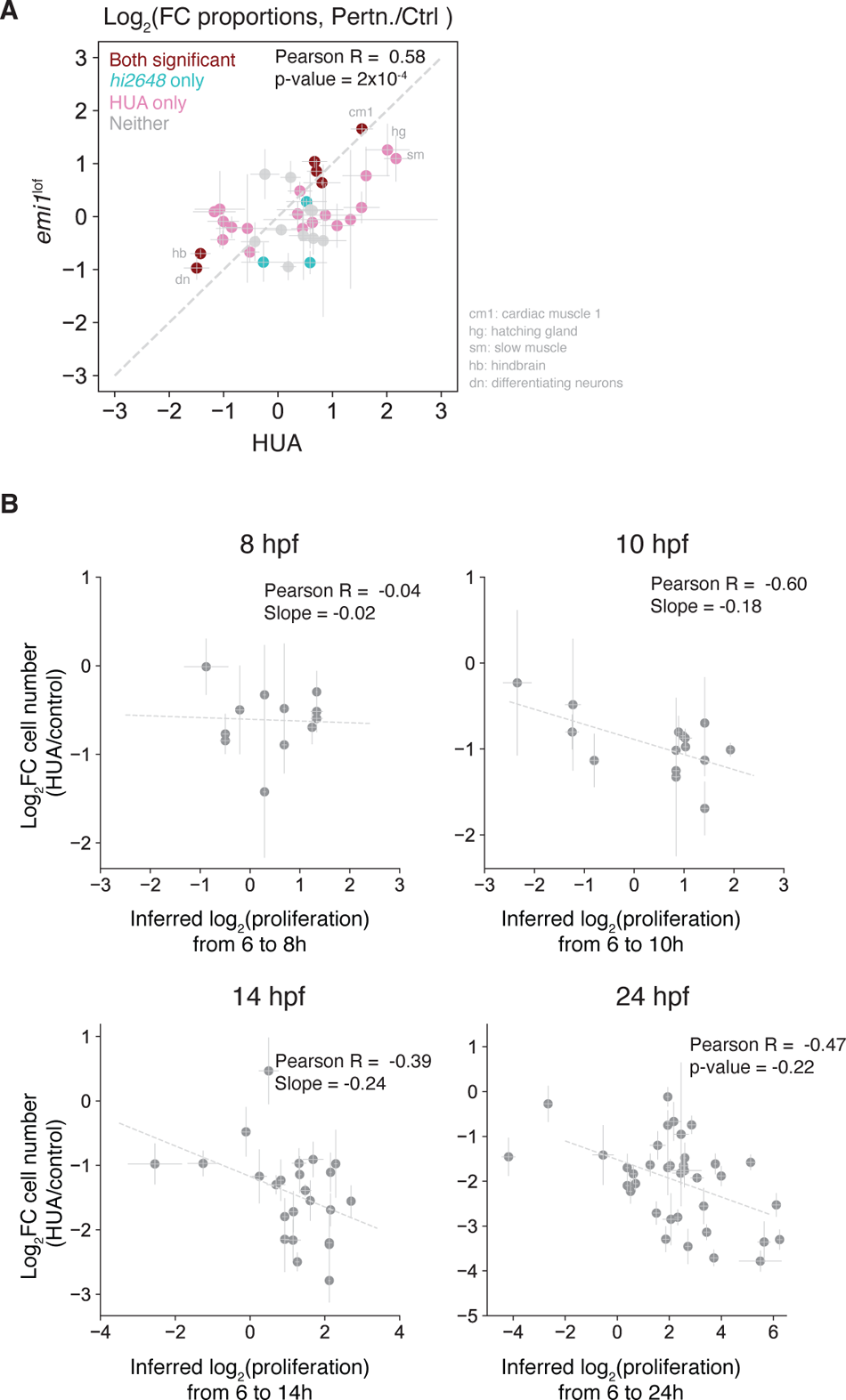
(A) Comparison between log_-_-fold-change in relative cell state abundances following cell cycle arrest with HUA treatment and *emi1* mutation. Each point is an annotated cell state. Brown: significant difference in fold change in both *emi1* mutant and HUA conditions with FDR < 0.05 using beta binomial test and Benjamini Hochberg correction; Teal: significant only in *emi1* mutant; Purple: significant only in HUA; Light grey: significant in neither condition. The comparison shows a high correlation in abundance changes between the two methods of cell cycle inhibition. (B) Comparison between log-fold-change in relative cell state abundances following cell cycle arrest by HUA and the expected proliferation lost during the period of cell cycle arrest (from 6hpf). Dashed grey line shows a linear fit for the data.

### Captions for supplementary tables

**Table S1**

Sheet 1: Assigned cell state names corresponding to the cell state labels in (*20*)

Sheet 2: Additional information regarding cell state classification.

**Table S2**

Number of cells in different conditions per 50k total cells as plotted in **Fig. 1G**

**Table S3**

Usages of cNMF programs in our dataset.

**Table S4**

Gene loadings for cNMF programs for 3000 highly variable genes.

**Table S5**

Top 50 genes enriched in the global perturbation associated programs.

**Table S6**

Fold changes in cell type proportions as plotted in **Fig. 5A**.

**Table S7**

Lineage relations between cell states at different time points in the published zebrafish development atlas (*20*). These were used to calculate proliferation score in **Fig. 5B-D**.

## References

1. F. Cremisi, A. Philpott, S. I. Ohnuma, Cell cycle and cell fate interactions in neural development. Curr Opin Neurobiol. 13, 26–33 (2003).

2. Y. Budirahardja, P. Gonczy, Coupling the cell cycle to development. Development (2009), doi:10.1242/dev.021931.

3. S. I. Ohnuma, A. Philpott, W. A. Harris, Cell cycle and cell fate in the nervous system. Curr Opin Neurobiol. 11, 66–73 (2001).

4. S. Dalton, Linking the Cell Cycle to Cell Fate Decisions. Trends Cell Biol. 25, 592–600 (2015).

5. R. Pop, J. R. Shearstone, Q. Shen, Y. Liu, K. Hallstrom, M. Koulnis, J. Gribnau, M. Socolovsky, A Key Commitment Step in Erythropoiesis Is Synchronized with the Cell Cycle Clock through Mutual Inhibition between PU.1 and S-Phase Progression. PLoS Biol. 8 (2010), doi:10.1371/journal.pbio.1000484.

6. H. Y. Kueh, Positive feedback between PU.1 and the cell cycle controls myeloid differentiation. Science. 342, 311 (2013).

7. F. Ali, C. Hindley, G. McDowell, R. Deibler, A. Jones, M. Kirschner, F. Guillemot, A. Philpott, Cell cycle-regulated multi-site phosphorylation of Neurogenin 2 coordinates cell cycling with differentiation during neurogenesis. Development. 138, 4267–4277 (2011).

8. A. E. Vernon, A. Philpott, A single cdk inhibitor, p27Xic1, functions beyond cell cycle regulation to promote muscle differentiation in Xenopus. Development. 130, 71–83 (2003).

9. K. Walsh, H. Perlman, Cell cycle exit upon myogenic differentiation. Curr Opin Genet Dev. 7, 597–602 (1997).

10. Z. Kadri, R. Shimizu, O. Ohneda, L. Maouche-Chretien, S. Gisselbrecht, M. Yamamoto, P. H. Romeo, P. Leboulch, S. Chretien, Direct binding of pRb/E2F-2 to GATA-1 regulates maturation and terminal cell division during erythropoiesis. PLoS Biol. 7 (2009), doi:10.1371/journal.pbio.1000123.

11. L. A. Buttitta, B. A. Edgar, Mechanisms controlling cell cycle exit upon terminal differentiation. Curr Opin Cell Biol. 19, 697–704 (2007).

12. P. H. Edgar, Bruce A, Lehman Dara A. and O’Farrell, Transcriptional regulation of string(cdc25): a link between developmental programming and the cell cycle. Development. 120, 3131–3143 (1994) (1994).

13. N. Satoh, On the clock mechanism determining the time of tissue-specific enzyme development during ascidian embryogenesis. J Embryol Exp Morphol. 54, 131–139 (1979).

14. V. Hartenstein, J. W. Posakony, Sensillum Development in the Absence of Cell Division: The Sensillum Phenotype of the Drosophila Mutant string. Dev Biol. 138, 147–158 (1989).

15. W. A. Harris, V. Hartenstein, Neuronal Determination in Xenopus Embryos without Cell Division. Cell. 6, 499–515 (1991).

16. B. B. Riley, E. M. Sweet, R. Heck, A. Evans, K. N. McFarland, R. M. Warga, D. A. Kane, Characterization of harpy/Rca1/emi1 mutants: Patterning in the absence of cell division. Developmental Dynamics. 239, 828–843 (2010).

17. L. Zhang, C. Kendrick, D. Julich, S. A. Holley, Cell cycle progression is required for zebrafish somite morphogenesis but not segmentation clock function. Development. 135, 2065–2070 (2008).

18. J. A. Farrell, Y. Wang, S. J. Riesenfeld, K. Shekhar, A. Regev, A. F. Schier, Single-cell reconstruction of developmental trajectories during zebrafish embryogenesis. *Science*, eaar3131 (2018).

19. X. Jin, S. K. Simmons, A. Guo, A. S. Shetty, M. Ko, L. Nguyen, V. Jokhi, E. Robinson, P. Oyler, N. Curry, G. Deangeli, S. Lodato, J. Z. Levin, A. Regev, F. Zhang, P. Arlotta, In vivo Perturb-Seq reveals neuronal and glial abnormalities associated with autism risk genes. Science 370 (2020)

20. D. E. Wagner, C. Weinreb, Z. M. Collins, J. A. Briggs, S. G. Megason, A. M. Klein, Single-cell mapping of gene expression landscapes and lineage in the zebrafish embryo. Science. 360, 981– 987 (2018).

21. C. B. Kimmel, W. W. Ballard, S. R. Kimmel, B. Ullmann, T. F. Schilling, Stages of embryonic development of the zebrafish. Developmental dynamic. 203, 253–310 (1995).

22. C. B. Kimmel, R. M. Warga, T. F. Schilling, Origin and organization of the zebrafish fate map. Development. 108, 581–594 (1990).

23. Y. J. Machida, A. Dutta, The APC/C inhibitor, Emi1, is essential for prevention of rereplication. Genes Dev. 21, 184–94 (2007).

24. J. D. R. Reimann, E. Freed, J. Y. Hsu, E. R. Kramer, J.-M. Peters, P. K. Jackson, Emi1 is a mitotic regulator that interacts with Cdc20 and inhibits the anaphase promoting complex. Cell. 105(**5**), 645–655 (2001).

25. B. K. Tusi, S. L. Wolock, C. Weinreb, Y. Hwang, D. Hidalgo, R. Zilionis, A. Waisman, J. R. Huh, A. M. Klein, M. Socolovsky, Population snapshots predict early haematopoietic and erythroid hierarchies. Nature Publishing Group. 555 (2018), doi:10.1038/nature25741.

26. D. Kotliar, A. Veres, M. A. Nagy, S. Tabrizi, E. Hodis, D. A. Melton, P. C. Sabeti, Identifying gene expression programs of cell-type identity and cellular activity with single-cell RNA-Seq. Elife. 8 (2019), doi:10.7554/eLife.43803.

## References

27. J. Rhodes, A. Amsterdam, T. Sanda, L. A. Moreau, K. McKenna, S. Heinrichs, N. J. Ganem, K. W. Ho, D. S. Neuberg, A. Johnston, Y. Ahn, J. L. Kutok, R. Hromas, J. Wray, C. Lee, C. Murphy, I. Radtke, J. R. Downing, M. D. Fleming, L. E. MacConaill, J. F. Amatruda, A. Gutierrez, I. Galinsky, R. M. Stone, E. A. Ross, D. S. Pellman, J. P. Kanki, A. T. Look, Emi1 Maintains Genomic Integrity during Zebrafish Embryogenesis and Cooperates with p53 in Tumor Suppression. https://doi.org/10.1128/MCB.00558-09. 29, 5911–5922 (2023).

28. W. A. Harris, V. Hartenstein, Neuronal determination without cell division in xenopus embryos. Neuron. 6, 499–515 (1991).

29. A. M. Klein, D. A. Weitz, M. W. Kirschner Correspondence, Droplet Barcoding for Single-Cell Transcriptomics Applied to Embryonic Stem Cells. Cell. 161, 1187–1201 (2015).

30. R. Zilionis, J. Nainys, A. Veres, V. Savova, D. Zemmour, A. M. Klein, L. Mazutis, Single-cell barcoding and sequencing using droplet microfluidics. Nature Publishing Group. 12 (2016), doi:10.1038/nprot.2016.154.

31. H. M. T. Choi, M. Schwarzkopf, M. E. Fornace, A. Acharya, G. Artavanis, J. Stegmaier, A. Cunha, N. A. Pierce, Third-generation in situ hybridization chain reaction: Multiplexed, quantitative, sensitive, versatile, robust. Development (Cambridge). 145 (2018), doi:10.1242/dev.165753.

32. K. J. Livak, T. D. Schmittgen, Analysis of Relative Gene Expression Data Using Real-Time Quantitative PCR and the 2−ΔΔCT Method. Methods. 25, 402–408 (2001).

33. K. Polański, M. D. Young, Z. Miao, K. B. Meyer, S. A. Teichmann, J. E. Park, BBKNN: fast batch alignment of single cell transcriptomes. Bioinformatics. 36, 964–965 (2020).

34. R. Zilionis, C. Engblom, C. Pfirschke, V. Savova, E. Levantini, M. J. Pittet, A. M. H. Klein, M. J. P.) Edu, Single-Cell Transcriptomics of Human and Mouse Lung Cancers Reveals Conserved Myeloid Populations across Individuals and Species. Immunity. 50, 1317–1334.e10 (2019).

35. T. V. Pham, C. R. Jimenez, An accurate paired sample test for count data. Bioinformatics. 28, i596–i602 (2012).

36. T. V. Pham, S. R. Piersma, M. Warmoes, C. R. Jimenez, On the beta-binomial model for analysis of spectral count data in label-free tandem mass spectrometry-based proteomics. Bioinformatics. 26, 363–369 (2010).

37. V. K. Mootha, C. M. Lindgren, K. F. Eriksson, A. Subramanian, S. Sihag, J. Lehar, P. Puigserver, E. Carlsson, M. Ridderstråle, E. Laurila, N. Houstis, M. J. Daly, N. Patterson, J. P. Mesirov, T. R. Golub, P. Tamayo, B. Spiegelman, E. S. Lander, J. N. Hirschhorn, D. Altshuler, L. C. Groop, PGC-1α-responsive genes involved in oxidative phosphorylation are coordinately downregulated in human diabetes. Nature Genetics 2003 34:3. **34**, 267–273 (2003).

38. A. Subramanian, P. Tamayo, V. K. Mootha, S. Mukherjee, B. L. Ebert, M. A. Gillette, A. Paulovich, S. L. Pomeroy, T. R. Golub, E. S. Lander, J. P. Mesirov, Gene set enrichment analysis: A knowledge-based approach for interpreting genome-wide expression profiles. Proc Natl Acad Sci U S A. 102, 15545–15550 (2005).

